# Regularized selection indices for breeding value prediction using hyper-spectral image data

**DOI:** 10.1101/625251

**Authors:** Marco Lopez-Cruz, Eric Olson, Gabriel Rovere, Jose Crossa, Susanne Dreisigacker, Suchismita Mondal, Ravi Singh, Gustavo de los Campos

**Author notes:** Correspondence and request should be addressed to G.D.L.C.

## Abstract

High-throughput phenotyping (HTP) technologies can produce data on thousands of phenotypes per unit being monitored. These data can be used to breed for economically and environmentally relevant traits (e.g., drought tolerance); however, incorporating high-dimensional phenotypes in genetic analyses and in breeding schemes poses important statistical and computational challenges. To address this problem, we developed regularized selection indices; the methodology integrates techniques commonly used in high-dimensional phenotypic regressions (including penalization and rank-reduction approaches) into the selection index (SI) framework. Using extensive data from CIMMYT’s (International Maize and Wheat Improvement Center) wheat breeding program we show that regularized SIs derived from hyper-spectral data offer consistently higher accuracy for grain yield than those achieved by canonical SIs, and by vegetation indices commonly used to predict agronomic traits. Regularized SIs offer an effective approach to leverage HTP data that is routinely generated in agriculture; the methodology can also be used to conduct genetic studies using high-dimensional phenotypes that are often collected in humans and model organisms including body images and whole-genome gene expression profiles.

## Introduction

High-throughput phenotyping (HTP) technologies have been adopted at a fast pace in agriculture; applications range from the use of HTP in highly controlled environments (e.g., growth chambers^1^) to extensive HTP using sensing devices mounted on aerial (e.g., hyper-spectral cameras mounted on aerial vehicles^2^) and terrestrial equipment such as tractors and combine harvesters^3^. Modern agricultural production systems use HTP data to optimize management practices^4^, forecast agricultural outputs^5^ and to assess the quality (e.g., protein content) of agricultural commodities^6^. HTP data can also be a valuable input for breeding programs. For instance, extensive HTP may enable an expansion of genetic testing that can lead to higher intensity of selection and faster genetic progress. Moreover, HTP data may offer opportunities to improve traits such as drought tolerance that are otherwise difficult to measure and breed for.

Sensors can generate data on hundreds or thousands of phenotypes per unit being monitored. For example, hyper-spectral cameras can generate reflectance of electromagnetic power at hundreds of wavelengths in the visible and infrared spectrum. These measurements can be considered as indicator phenotypes that can be used to predict other traits. An extensive body of research deals with the use HTP data to predict phenotypes such as grain yield^5,7–9^, dry matter^3^, oil and protein content^10,11^. However, there has been much less research on how to integrate HTP data in genetic studies and in breeding schemes. In genetics, the problem of predicting the genetic merit of a target trait given a set of correlated phenotypes was first addressed by Smith^12^ and Hazel^13^ who introduced the concept of selection index (SI) in plant and animal breeding, respectively.

A SI seeks to improve a target trait *y*_*i*_ (e.g., grain yield) using information from another set of measured traits (e.g., hyper-spectral image data). A linear SI is a weighted sum of the measured phenotypes with weights derived to maximize the correlation between the genetic merit for the selection target and the SI. Thus, the SI methodology offers a natural framework for integrating HTP data into breeding decisions. However, when the measured phenotype is high-dimensional, the naïve application of the SI can lead to overfitting and sub-optimal accuracy of indirect selection.

To address this problem, we developed regularized selection indices (including penalized and reduced-rank methods) that are tailored to achieve accurate prediction of genetic values using high-dimensional phenotypes. The proposed methodology integrates into the SI framework methods often used to prevent overfitting in high-dimensional phenotypic regressions^14^. Using extensive multi-environment crop imaging data from CIMMYT’s wheat breeding program we show that regularized SIs offer improved accuracy of indirect selection in both optimal and stress environments.

## Results

A **selection index** is a linear combination of *p* measured phenotypes, ***x***_*i*_ = (*x*_*i*1_, …, *x*_*ip*_)′, of the form 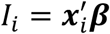, where ***β*** = (*β*_1_, …, *β*_*p*_)′ is a vector of regression coefficients whose entries define the weights of each of the measured phenotypes in the SI. In a canonical SI those weights are derived by minimizing the expected squared deviation between the genetic merit for the selection goal (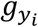, e.g., the genetic merit for grain yield of the *i*^th^ genotype) and the index, that is:

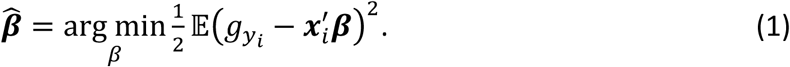

The solution to this optimization problem is (see *Methods* section):

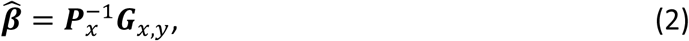

where 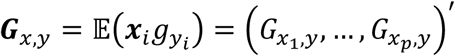 is a vector containing the genetic covariances between the selection objective (*y*_*i*_) and each of the measured traits (***x***_*i*_), and ***P***_*x*_ is the (population) phenotypic variance-covariance matrix of the measured phenotypes, that is, 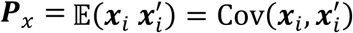. Thus, a canonical SI takes the form 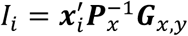. The theory underlying the derivation of SIs and response to indirect selection is well established^15,16^.

The SI is by construction the best linear predictor (BLP) of the genetic merit for the selection goal; this property holds when ***G***_*x,y*_ and ***P***_*x*_ are known. However, when the number of measured phenotypes is large, errors in the estimation of ***P***_*x*_ and ***G***_*x,y*_ may lead to overfitting and sub-optimal accuracy of indirect selection.

### Regularized selection indices

Reduced-rank (e.g., principal components methods) and penalized regression^14^ are two approaches commonly used to confront overfitting in high-dimensional regression problems. These methodologies were developed for regression problems involving an observable phenotype (*y*_*i*_). In the SI, the response 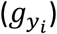 is unobservable; however, the same principles that are applied in phenotypic reduced-rank and penalized regressions can be integrated into the SI framework.

### Reduced-rank selection indices

In principal components (PC) regression, the response is regressed on a reduced number (*q* < *p*) of PCs extracted from a set of predictors (***x***_*i*_); the same concept can be used to derive a reduced-rank SI. For instance, one can extract a reduced number of PCs from a crop image and the resulting PCs can be used as ‘measured traits’ in equation (1).

The solution of equation (1) will render estimates of the regression coefficients for the PCs, which can be transformed back to coefficients applicable to the measured traits (see *Methods*). Thus, a reduced-rank SI (referred to as PC-SI) can be derived following these steps: (*i*) extract, using the singular value decomposition, *q* PCs from the matrix containing the measured phenotypes, (*ii*) estimate the genetic covariances between the first *q* PCs and the selection objective, (*iii*) use these estimated (co)variances to derive coefficients associated with the top *q* PCs; finally, (*iv*) transform these coefficients into coefficients for the measured phenotypes. This process can be done using *q* = 1,2, …, *p* PCs (*q* = *p* renders the canonical SI). For the sequence of estimates 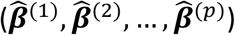, one can evaluate the accuracy of indirect selection of the resulting SI and an *optimal rank* for the PC-SI can be chosen to maximize the accuracy of indirect selection.

### Penalized selection indices

In a penalized regression, regularization is achieved by including in the objective function a penalty on model complexity. In the context of a SI, we have

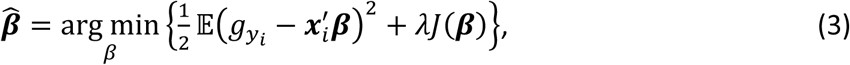

where *λ* is a penalty parameter (*λ* = 0 yields the coefficients for the canonical SI) and *J*(***β***) is a penalty function. Commonly used penalties include the L2 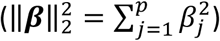 and L1 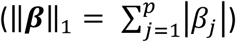 norms^17^, or a weighted sum of the two^18^.

Using 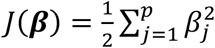 in equation (3) renders a **Ridge-regression-type PSI** (RR-PSI, see *Methods*):

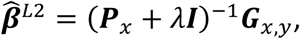

where ***I*** is a *p* × *p* identity matrix. The RR-PSI (referred to as the L2-PSI) yields shrunken estimates of the regression coefficients.

In many applications, variable selection (i.e., a SI that is a function of a subset of the measured phenotypes) may be desirable. This property can be obtained using penalties involving the L1-norm, either alone, 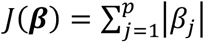 (LASSO^19^), or in combination with the L2-norm, 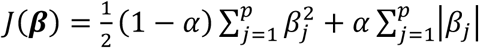 (elastic-net^18^). Unlike the L2-PSI, the LASSO and elastic-net SIs (hereinafter referred to as L1-PSI and EN-PSI, respectively) do not have a closed-form solution^14^. However, solutions for PSIs involving an L1-penalty can be obtained using iterative procedures such as the coordinate descent^20^ and the least angle regression^21^ (LARS) algorithms (see *Methods*). As with the PC-SI, an optimal PSI can be obtained by choosing the values of the regularizing parameters (*λ, α*) that maximize the accuracy of indirect selection.

### Accuracy of indirect selection

Indirect selection accuracy is defined as the correlation between the index used to rank genotypes and the genetic merit of the selection objective, that is, 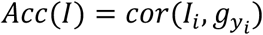. This parameter is equal to the product of the square root of the heritability of the SI (*h*_*I*_) times the genetic correlation between the SI and the selection target, 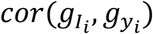^16^. To avoid estimation bias *Acc*(*I*) must be estimated using data that was not used to derive the coefficients of the index (Fig. 1); therefore, in the application presented below we: (*i*) partitioned the data into training and testing sets, (*ii*) derived the coefficients of the SI in the training set, (*iii*) applied these coefficients to image data of the testing set 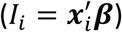, and (*iv*) estimated 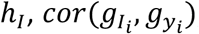, and *Acc*(*I*) in the testing set. Furthermore, we quantified the efficiency of indirect selection relative to mass phenotypic selection (RE) using 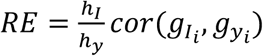^16^.

**Figure 1.**
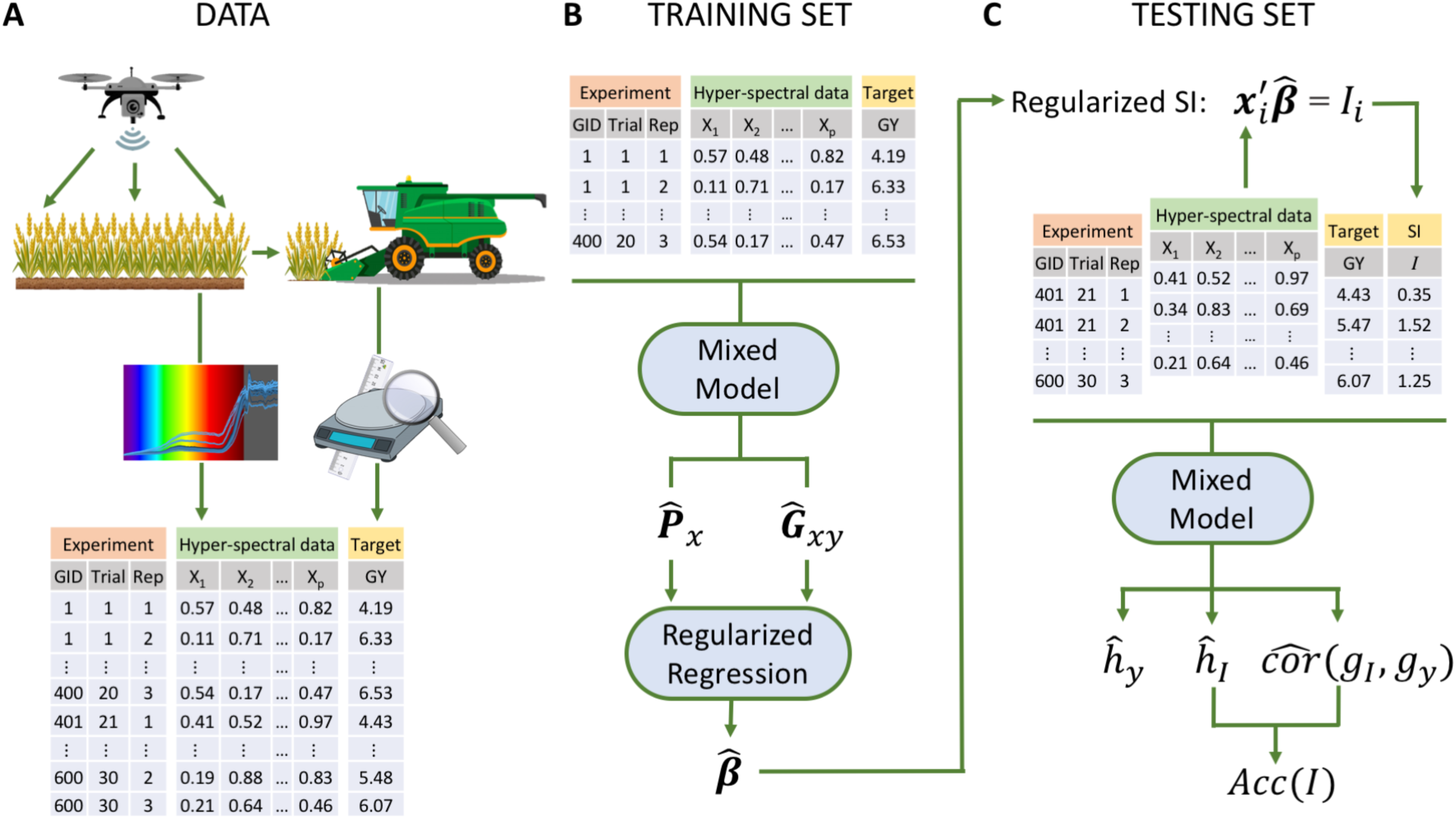
Prediction of the genetic merit for grain yield using hyper-spectral crop image data. (A) Data consists of hyper-spectral reflectance data (***x***_*i*_) and phenotypic measurements of the target trait (*y*_*i*_, e.g., grain yield). (B) A subset of the data (the training set) is used to derive the coefficients (***β***) of a selection index. (C) These coefficients are then applied to image data of individuals in the testing set to derive the index (*I*_*i*_) for each individual. The predictive ability of the index is assessed by calculating the accuracy of indirect selection (*Acc*(*I*)) in the testing set.

### Regularized selection indices for wheat grain yield using hyper-spectral image data

We applied the methodology described in the previous section to data (*n*=3,276) from the CIMMYT’s Global Wheat Program consisting of grain yield (ton ha^−1^) and hyper-spectral image data. The data were collected at CIMMYT’s experimental station in Ciudad Obregon, Sonora, Mexico (27°20′ N, 109°54′ W, 38 masl) from 39 yield trials in which a total of 1,092 genotypes were tested. Rainfall in Obregon is very limited; therefore, four different environments were generated representing a combination of planting methods (*Flat* or *Bed*), controlled irrigation (minimal, 2 or 5 irrigations), and planting dates (optimum or early-heat). As expected, average yield decreased as drought stress intensity increased (see Table 1 and Supplementary Fig. S1 for boxplots of yield by environment).

**Table 1.** Average grain yield and heritability by environmental condition

| Planting conditions |  | Number of irrigations | Abbreviation | Average (SD) Yield | Heritability (SD) |
| --- | --- | --- | --- | --- | --- |
| Date | System |  |  |  |  |
| Optimum | Flat | Minimal | Flat-Drought | 2.06 (0.58) | 0.83 (0.016) |
|  | Bed | 2 | Bed-2IR | 3.67 (0.43) | 0.66 (0.032) |
|  |  | 5 | Bed-5IR | 6.11 (0.61) | 0.43 (0.025) |
| Early |  | 5 | Bed-EHeat | 6.43 (0.73) | 0.61 (0.018) |

SD: standard deviation.

Image data was collected using an infrared and an hyper-spectral camera and consisted of reflectance of electromagnetic power at 250 wavebands within the visible and near-infrared spectrums (392-850 nm). Separate images were collected at 9 time-points covering vegetative (VEG), grain filling (GF), and maturity (MAT) stages of the crop (see Supplementary Fig. S2). Grain yield and image data were pre-adjusted using mixed-effects model that accounted for genotype, trial, replicate, and sub-block (see *Methods* section).

### Regularization improves the heritability and the accuracy of the index

To assess the effect of regularization on the accuracy of indirect selection we fitted an L1-PSI over a grid of values of the regularization parameter (*λ*^(1)^ > *λ*^(2)^ > … > 0 in equation (3), using *λ* = 0 renders a canonical SI). For each of the solutions 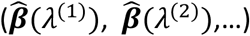 we estimated the heritability of the resulting index and the genetic correlation between the index and the selection target, and from those estimates we derived the accuracy of indirect selection. The same approach was used to evaluate the accuracy of indirect selection of PC-SIs with a varying number (1, 2, …) of PCs.

We first fitted PSIs and PC-SIs using data from a single time-point; the results from the latest time-point (corresponding to MAT or late GF stages depending on the environment) are presented in Fig. 2 (see Supplementary Figs. S3-S5 for other time-points). The heritability of the L1-PSI (Fig. 2A) decreased as more bands became active in the index. Likewise, the heritability of PC-SI (Fig. 2B) decreased with the number of PCs used. However, the genetic correlation increased as either more bands become active in the L1-PSI or more PCs were used in the PC-SI. Consequently, the maximum accuracy of indirect selection was achieved with a SI of intermediate complexity (with anywhere between 20 and 60 of the 250 bands being active in the L1-PSI, and between 20-60 PCs in the PC-SI). Results for other time-points and environments (Supplementary Figs. S3-S5) exhibited similar patterns with some differences between environments. The accuracy of indirect selection of the optimal L1-PSI was always close to that of the optimal PC-SI and that of the optimal L2-PSI (Supplementary Table S1). Importantly, in all cases the accuracy of indirect selection of the optimal regularized SIs was considerably higher than that of the canonical SI, which is the one corresponding to 250 active bands or 250 PCs (i.e., the right-most results in the plots in Fig. 2).

**Figure 2.**
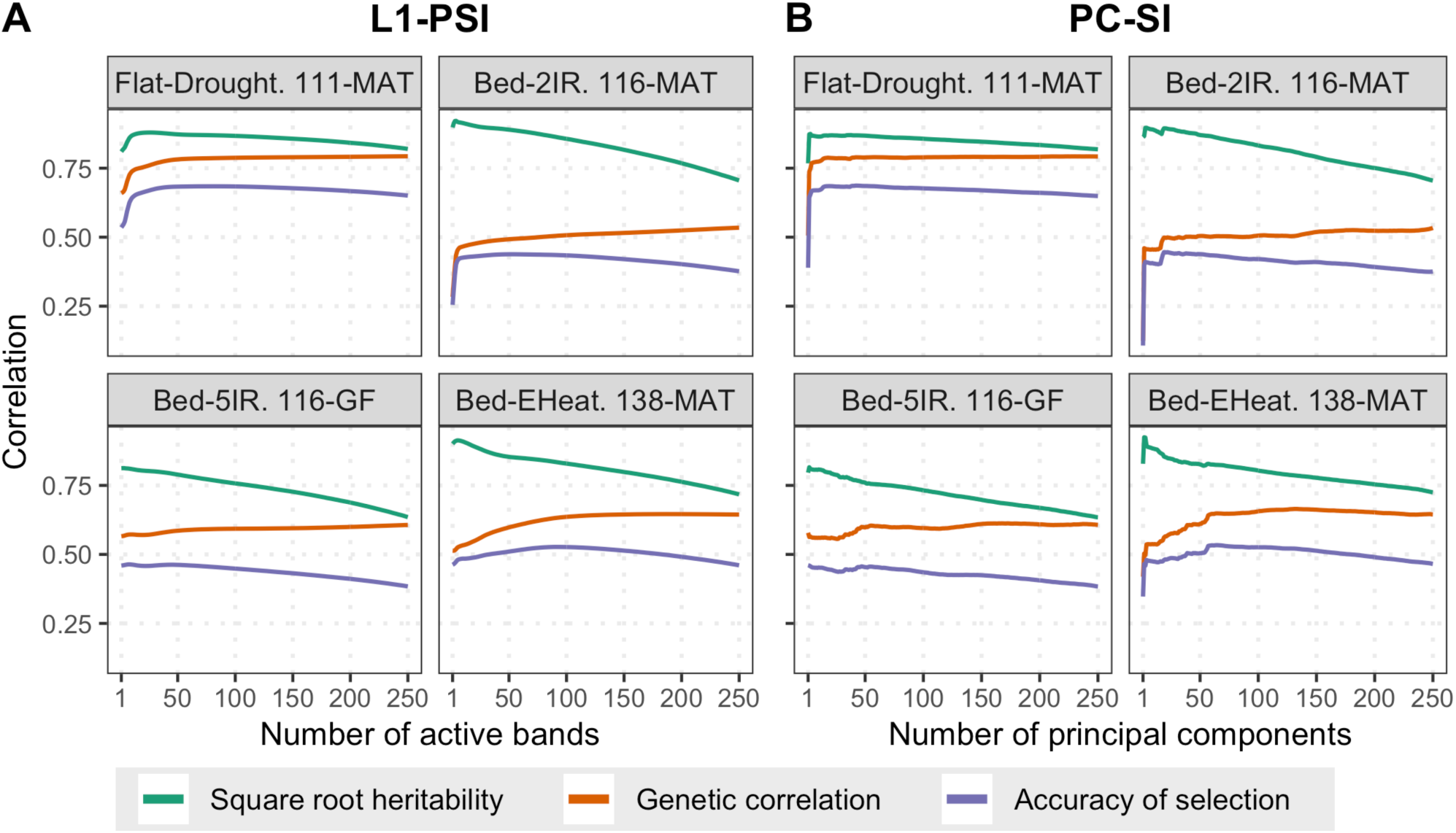
Accuracy of indirect selection of regularized SIs and its components. Square root heritability (green), genetic correlation (orange), and accuracy of indirect selection (purple, all averaged over 100 training-testing partitions), versus the number of predictors used to build the index: (A) number of active bands in the case of the L1-PSI, or (B) number of PCs in the PC-SI. Each panel represents one environment (latest time-point).

Fig. 3 displays the accuracy of indirect selection across time-points for the optimal (i.e., the one with the highest accuracy of indirect selection) L1-PSI and PC-SI. For comparison we also display in the plot the accuracy of indirect selection achieved by a canonical SI (in green). The estimated 95% confidence intervals of the accuracy of the regularized SIs (either PC-SI or L1-PSI) are all above (and do not overlap) with the confidence intervals for the accuracy of the canonical SI, except for a single time-point (57 DAS in environment *Bed-2IR*). Results from Tukey’s Honest Significance Difference confirmed that the accuracy of the regularized SIs is statistically different (higher) than the canonical SI at a 5% of significance (Supplementary Table S1) for all but one (57 DAS in environment *Bed-2IR*) time-point/environment. Regularization increased the selection accuracy across time-points and environments. Regularized SIs (either PC-SI or L1-PSI) had an accuracy of indirect selection that was in average 10-40% higher than the accuracy achieved by a canonical SI. These gains in accuracy were stronger in the optimal environment (*Bed-5IR* with a median of 36%) and smaller in the stressed environments (*Flat-Drought* and *Bed-EHeat* with a median of 16%). Interestingly, there were no sizable differences between the accuracy of indirect selection achieved with the optimal L1-PSI and that of the optimal PC-SI. Compared with the canonical SI, regularized SIs had higher heritability (Supplementary Fig. S6); this was achieved without compromising the genetic correlation (Supplementary Fig. S7), thus leading to a higher accuracy of indirect selection achieved by either penalization or rank-reduction strategies.

**Figure 3.**
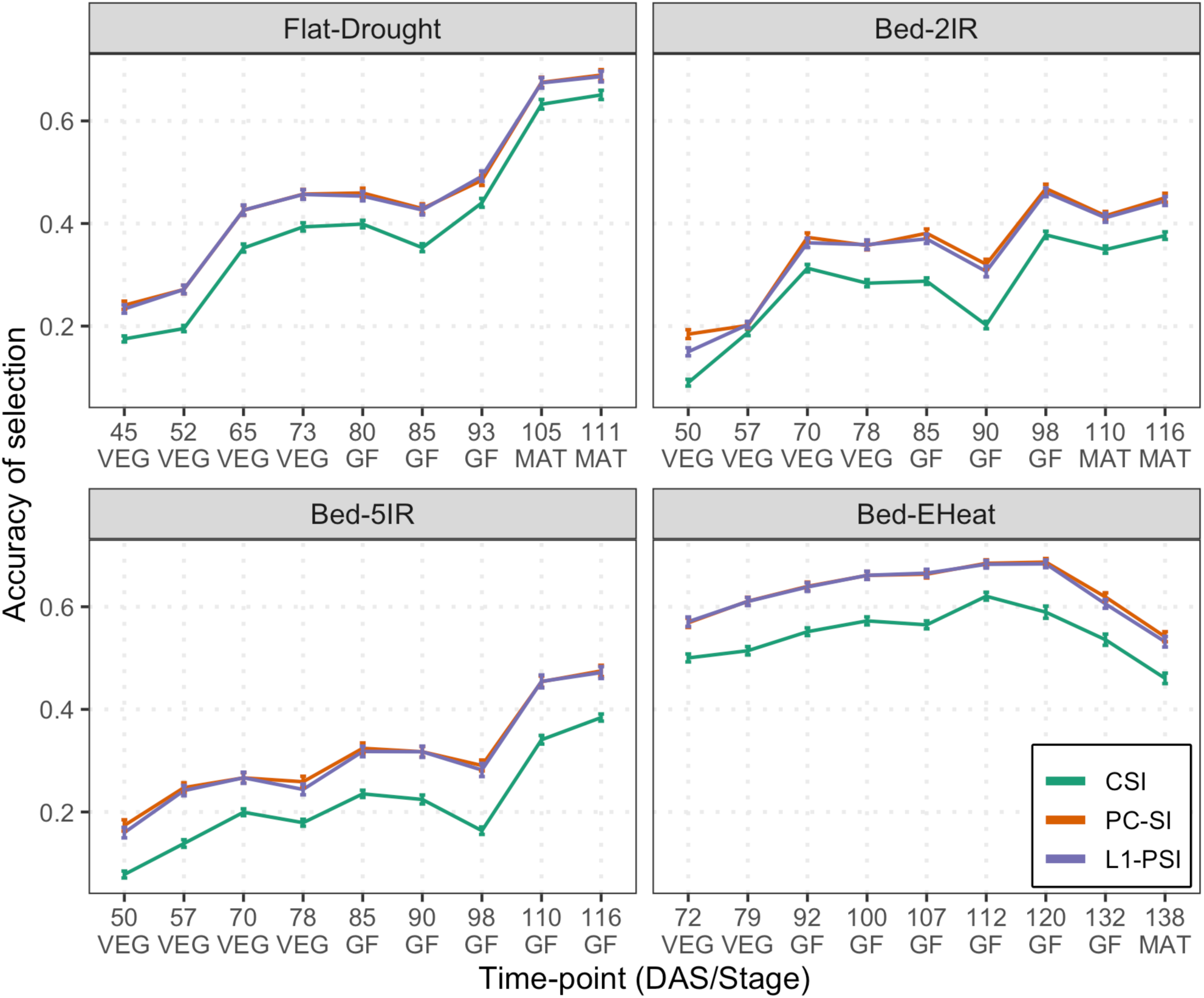
Accuracy of indirect selection achieved by a canonical (CSI) and by regularized (PC-SI and L1-PSI) selection indices. The lines provide the average accuracy over 100 training-testing partitions. Vertical lines represent a 95% confidence interval for the average. The horizontal axis give the time-point at which images were collected and are expressed in both days after sowing (DAS) and stages (VEG=vegetative, GF=grain filling, MAT=maturity).

### Using data from multiple time-points further improves selection accuracy

The results presented above were based on data from a single time-point. We also generated selection indices using data from multiple time-points (in this case, ***x***_*i*_ was a vector containing 2,250 traits, corresponding to 250 wavebands measured at each of 9 time-points). Integrating data from multiple time-points further increased the accuracy of L1-PSI by a margin that ranged from 1 to 8 points on the correlation scale (Table 2). The gains in selection accuracy obtained using data from multiple time-points were more evident in environments with lower accuracy; similar results were obtained for the PC-SI and L2-PSI (Supplementary Table S1).

**Table 2.**
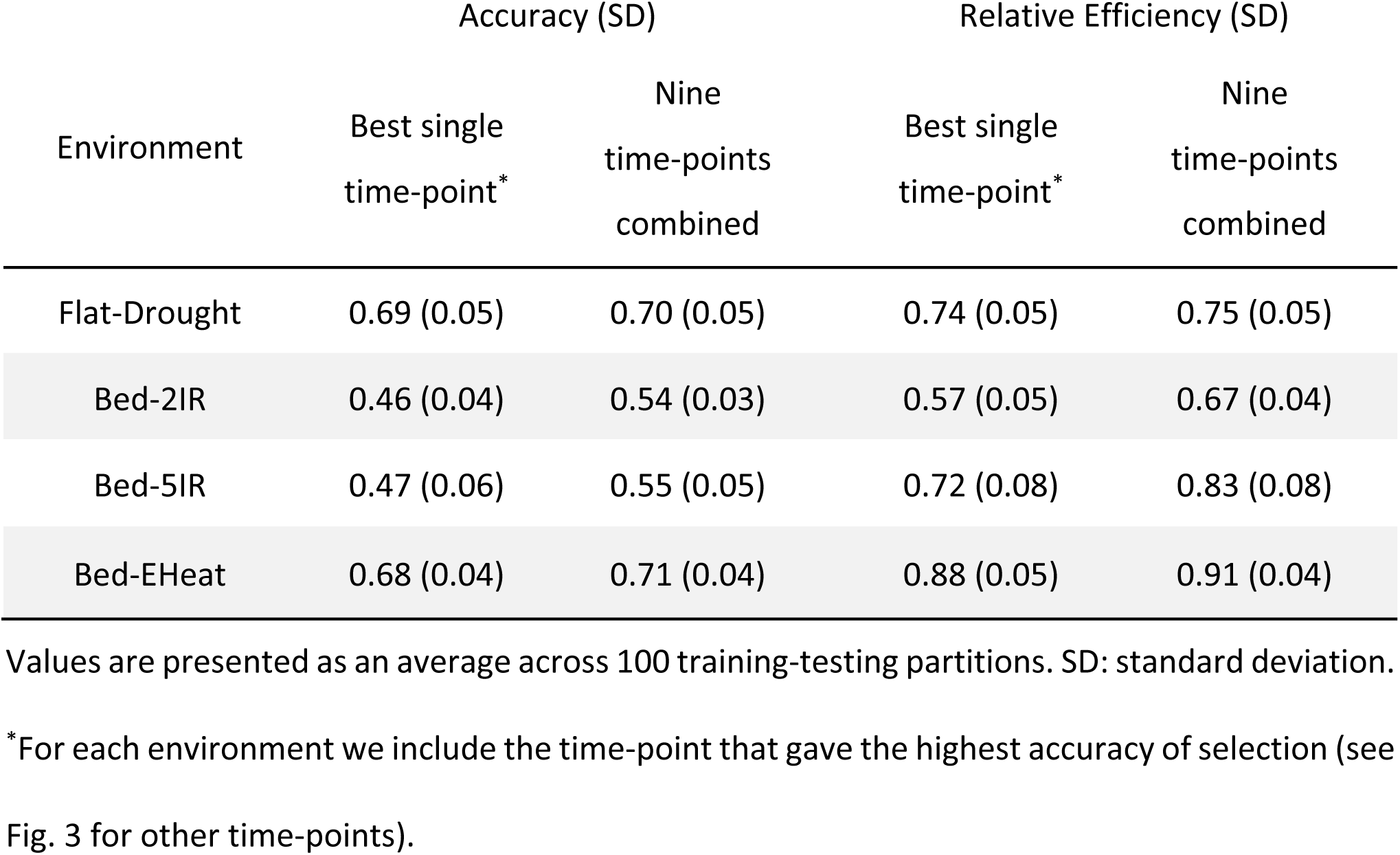
Accuracy and relative efficiency of indirect selection of an L1-penalized SI using data from one and nine time-points.

### L1-penalization leads to sparse selection indices

Fig. 4 shows a heatmap for the solutions of the optimal L1-PSI that integrated data from the 9 time-points. Each panel represents an environment, horizontal bands represent time-points. Within each time-point wavebands not entering in the solution are in grey and non-zero coefficients are represented in a yellow-red scale (red indicates large absolute-value coefficients). The well-irrigated environments (*Bed-5IR* and *Bed-EHeat*) had considerably sparser indices with only a reduced number of wavebands in the solutions; these were mostly located in the violet, blue and red regions of the spectrum. In stressed environments (*Flat-Drought* and *Bed-2IR*) there were also a few wavebands in the green and infrared regions that were active. In all the indices, there were wavebands from several time-points that were active in the optimal solution, suggesting that data from both early and late phenological stages are informative about the genetic merit for grain yield.

**Figure 4.**
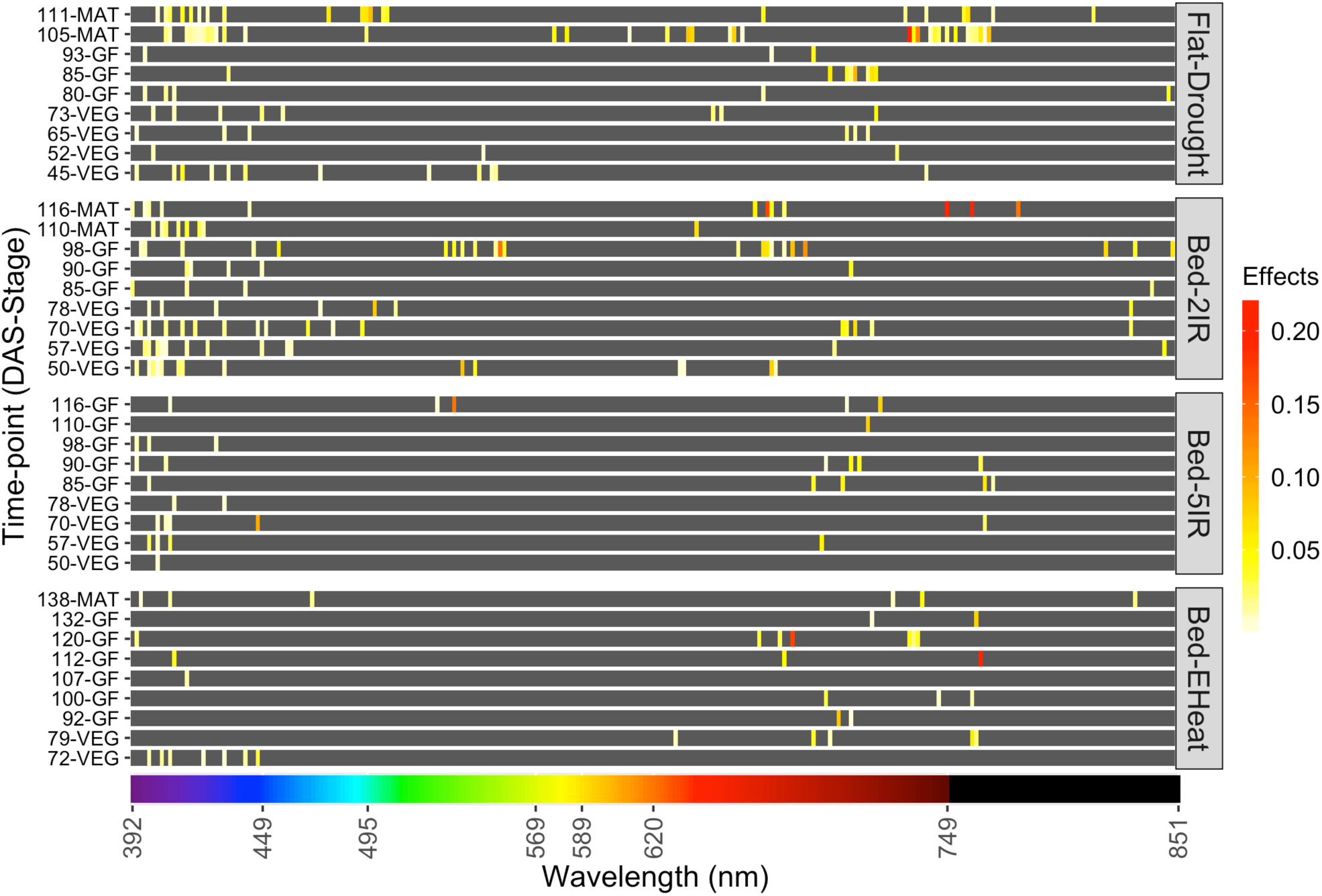
Heatmap of regression coefficients for L1-penalized selection indices. Separate indices were derived for each environment using multi time-point data. DAS=days after sowing, VEG, GF, MAT represent vegetative, grain-filling and maturity stages, respectively. The bottom color-bar shows the light color associated with each waveband in the visible spectrum (≤ 750 nm); black was used to represent the near-infrared spectrum (wavelength > 750 nm).

### Comparison with phenotypic prediction

We compared the accuracy of indirect selection of the PSI and PC-SI with vegetation indices and penalized phenotypic prediction. Vegetation indices are often used to predict yield^22^, biomass, and chlorophyll content^23,24^. We considered two vegetation indices: the Red and Green Normalized Difference Vegetation Indices (RNDVI^25^ and GNDVI^26^ respectively). For each of these indices we estimated the genetic correlation with grain yield, as well as their heritability and accuracy of indirect selection (Supplementary Table S1). Overall, the accuracy of indirect selection of the GNDVI and RNDVI was lower than the one achieved with a PSI (the average difference in accuracy between RNDVI and the L1-PSI varied by environment from 0.02 to 0.14 points in correlation, Supplementary Table S1, in favor of the L1-PSI). The heritability of the GNDVI and RNDVI was similar and superior in some cases to that of the L1-PSI (Supplementary Fig. S6); however, the genetic correlation between the vegetation indices and grain yield was (in most time-points and environments) lower than the genetic correlation between the L1-PSI and grain yield (Supplementary Fig. S7). Thus, the main driver of the difference in accuracy between the L1-PSI and the vegetation indices was the difference in genetic correlation.

We also fitted L1-penalized phenotypic prediction (L1-Phen) and compared the accuracy of indirect selection of these phenotypic prediction methods with that of penalized SIs. Overall, the L1-Phen achieved an accuracy of indirect selection very close to that of the L1-PSI (Supplementary Table S1); however, in a few environments at some time-points, the L1-PSI achieved a higher accuracy of indirect selection than the phenotypic prediction.

## Discussion

High-throughput phenotyping has been extensively adopted in agricultural research and commercial production. Extracting interpretable information from HTP data poses important statistical challenges. The clear majority of research in this area has focused on calibrating equations to predict phenotypes (e.g., total biomass, grain yield) using HTP data as inputs. This approach is well-suited for phenotypic prediction; however, the same approach can be sub-optimal for selection because the best predictor of a phenotype is not always the best predictor of the genetic merit of the same trait.

The best (linear) phenotypic predictor is the sum of the best linear predictor of the genetic merit (*g*_*n*_) plus the best linear predictor of the environmental term (*ε*_*n*_), that is, 𝔼(*y*|***x***) = 𝔼(*g*_*n*_|***x***) + 𝔼(*ε*_*n*_|***x***). The first term, 𝔼(*g*_*n*_|***x***), is the SI and it is, by construction, maximally correlated with the genetic merit. The second term, 𝔼(*ε*_*n*_|***x***), is relevant for phenotypic prediction but represents noise when the problem is that of selecting the best genotypes.

Selection indices exploit genetic covariances, while phenotypic prediction relies on phenotypic covariances between the selection target and the measured phenotype (e.g., crop imaging). Thus, the two methods yield different results whenever the patterns of phenotypic correlations are sufficiently different from the patterns of genetic correlations. In our data set, environmental conditions were highly controlled, with relatively low un-controlled within-trial variability in environmental conditions. Consequently, the patterns of phenotypic and genetic correlations were very similar (see Supplementary Fig. S8). This was true for many time-points and environments but not in others (e.g., 80, 85 and 93 DAS in *Flat-Drought*, and 90 and 98 DAS in *Bed-2IR*); it was exactly in those time-points and environments that the L1-PSI achieved higher accuracy of indirect selection than the L1-Phen method (Supplementary Table S1).

A canonical SI (equation (1)) is, by construction, maximally correlated with the genetic merit of the selection objective. This optimality property holds when the genetic and phenotypic (co)variance matrices that are needed to derive the coefficients of the SI (see equation (2)) are known without error. However, when the measured phenotype is high-dimensional, estimation errors in the phenotypic (co)variance matrix (***P***_*x*_), as well as in the genetic covariances (***G***_*x,y*_), can make the canonical SI sub-optimal. Our empirical results confirm this: canonical SIs over-fitted the data; this leads to a SI with low heritability and low accuracy of indirect selection.

To prevent overfitting, we considered integrating ideas commonly used in high-dimensional regression into the SI methodology. Our empirical results show that regularization consistently improves the accuracy of indirect selection relative to canonical SIs. We verified this for various environmental conditions and for crop imaging data collected at 9 different time-points. The optimal PSI and the optimal PC-SI achieved almost the same accuracy of indirect selection for all the environments and time-points, suggesting that either type of regularization can be effective.

Reduced-rank selection indices are appealing because after dimension reduction the problem of deriving a SI is trivial and can be dealt with methods commonly used to derive canonical SIs. Moreover, after HTP has been reduced to a few derived-traits (say the top 10 PCs), these traits can be integrated into genetic evaluations (either pedigree-based^27^ or genomic-enabled^28^) using standard multi-trait models.

Principal components-based methods have been considered before in the analysis of Fourier-transformed infrared (FTIR) spectra derived from milk samples. For instance, Soyeurt, Misztal & Gengler^29^ used a reduced number of FTIR-derived PCs to estimate variance components for selection objectives (e.g., fat or protein content in milk). Building upon this idea, Dagnachew, Meuwissen & Ådnøy^30^ suggested predicting the genetic merit for milk fatty acids using FTIR-derived PCs as ‘traits’ in a genetic evaluation. However, when mapping from genetic predictions of PC-lodgings onto genetic predictions for the selection objective the authors used coefficients derived from a phenotypic (partial least squares) regression. This does not guarantee that the resulting index is maximally correlated with the genetic merit of the selection target. The penalized and PC-SI presented in this study address that problem by using coefficients that are derived using genetic (and not phenotypic) covariances.

A disadvantage of the PC-SI is that the methodology does not naturally provide variable selection, a feature that may be desirable when the measured phenotype is high-dimensional.

Penalized selection indices can perform variable selection based on genetic covariances. While the derivation of a PSI is a bit more challenging than that of the PC-SI, the computational burden involved in the derivation of a PSI is not extremely high.

### Integration of PSI and PC-SI into genetic evaluations

The SIs considered here predict genetic merit for a selection target from a set of traits measured on an individual 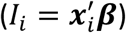; such indices exploit borrowing of information between traits within an individual. Borrowing of information between individuals increases selection accuracy; we envision two ways in which regularized SIs can be integrated into pedigree or genomic-based genetic evaluations.

One possibility is to use **a two-steps approach** whereas in the first step a PSI or a PC-SI is used to predict the genetic merit using within-individual information. This step can be considered as a task where patterns attributable to genetic covariances are extracted and those attributable to environmental covariances are smoothed-out. Then, in a second step, the resulting index-data {*I*_1_, …, *I*_*n*_} could be used as a trait in a genetic evaluation.

A **one-step** approach is also conceptually possible: the optimization problem of equation (3) can be modified by replacing ***x***_*i*_, the vector with the measured phenotypes on the *i*^th^ individual, with a vector 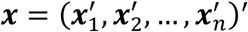 that contains all the available HTP data (measured on all *n* individuals); after expanding the squared error loss and taking expectations we get

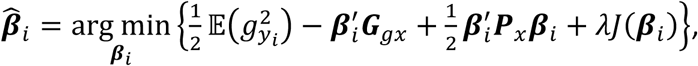

where ***G***_*gx*_ is a *pn* × 1 vector of genetic covariances including between-traits-within-individual (co)variances and between-subjects covariances. In standard genetic models, ***G***_*gx*_ takes a Kronecker form ***G***_*gx*_ = ***A***_*i*_ ∘ ***G***_*x,y*_, where ***A***_*i*_ are genetic (either DNA- or pedigree-derived) relationships between the candidate for selection and each of the individuals in the training set, and ***G***_*x,y*_ is, as before, a vector of genetic covariances between the selection objective and the measured traits (***x***). Likewise, ***P***_*x*_ is a *pn* × *pn* phenotypic (co)variance matrix. Estimating ***P***_*x*_ would require estimating all the genetic and environmental covariances among the measured traits. Therefore, while a one-step approach is conceptually feasible, the implementation can be computationally challenging.

### Impact of the use of high-throughput phenotypes in breeding programs

According to breeders’ equation^16,31^, the rate of genetic gain from selection is directly proportional to selection accuracy and selection intensity. Thus, relative to the use of canonical SIs, the use of regularized SIs is expected to increase selection gains by a factor equal to the gains observed in accuracy, that is between 10% and 40%. Relative to mass phenotypic selection, the PSIs had efficiencies, RE, ranging from 60% to 90%; therefore, relative to direct phenotypic selection, selection decisions based on PSI derived from images are expected to yield lower genetic gains than the ones that could be achieved via direct mass selection. However, the use of HTP technologies (e.g., crop monitoring using hyper-spectral cameras mounted in drones) may enable the expansion of the number of genotypes tested/measured as well as the number of locations where those genotypes are tested. This could lead to an increase in selection intensity which will in turn increase selection gains. For instance, if the use of HTP enables doubling the number of genotypes tested, the increase in selection gains that could be achieved with HTP may range from 20% (in the case where the PSI has RE of 60%) to 80% (for the traits/environments with RE of 90%).

### Regularized selection indices can also be a valuable tool in genetic research

High-dimensional phenotypes are also becoming increasingly available in genetic studies involving human subjects and model organisms. Performing genetic studies (e.g., genome-wide association analyses) on high-dimensional phenotypes is challenging and the burden of multiple testing across hundreds or thousands of phenotypes (e.g., RNA-abundance across thousands of genes) may critically compromise power. The PSI and PC-SI presented in this study could be used to extract genetic patterns from high dimensional phenotype data such as brain imaging or whole-genome gene expression profiles and these patterns can then be used as traits in genetic studies.

### Conclusion

We proposed two novel methods for predicting the genetic merit for selection objectives from high-dimensional phenotypes. These phenotypes are becoming increasingly available as the adoption of HTP in crop and animal production increases. The proposed methods integrate regularization procedures commonly used in high-dimensional regressions into the SI methodology. Regularization prevents overfitting and increases the accuracy of the index. The methods proposed here can be used to extract genetic patterns from almost any kind of high-dimensional phenotype, including not only HTP data emerging in agriculture but also high-dimensional phenotypes that emerge in genetic studies involving human subjects and model organisms.

## Methods

### Canonical selection index

The weights on a SI are derived as the solution to the optimization problem of equation (1):

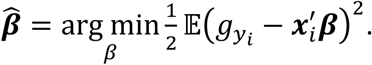

The right-hand side can be expressed as 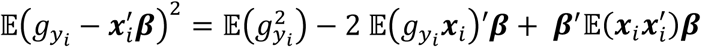. The first term, 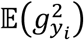, does not involve ***β***; therefore, it can be dropped from the objective function. Furthermore, if ***x***_*i*_ has null mean, and assuming that the environmental effects on ***x***_*i*_ are orthogonal to 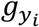, then 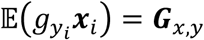 is a vector containing the genetic covariances between the selection target and each of the measured phenotypes. Likewise, 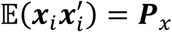 is the phenotypic (co)variance matrix of ***x***_*i*_. Therefore, the problem in equation (1) can be written as

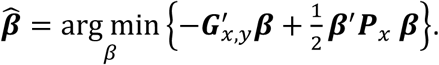

Differentiating the right-hand side with respect to vector ***β*** and setting the derivatives equal to zero leads to the first order conditions: 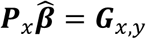; therefore,

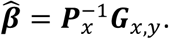

### Reduced-rank selection index

Recall that the singular value decomposition of a real-valued matrix, ***X*** = [***x***_1_, ***x***_2_, …, ***x***_*n*_]′ (individuals in rows, phenotypes in columns) takes the form ***X*** = ***UDV***′, where ***U*** = [***u***_1_, …, ***u***_*p*_] is the matrix containing the left-singular vectors that span the row-space of ***X, V*** = [***v***_1_, …, ***v***_*p*_] is the matrix with the right-singular vectors, and ***D*** = diag(*d*_1_, …, *d*_*p*_) is a diagonal matrix with positive or zero elements. The PCs ***W*** = ***XV*** = ***UD*** are linear combinations of the measured phenotypes. A reduced-rank regression uses the first *q* PCs 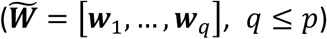 as ‘measured phenotypes’ in the SI:

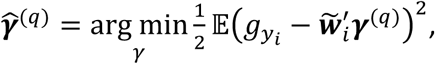

where 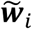 is a vector containing the scores for the *i*^th^ observation on the first *q* PCs. The solution to the optimization problem takes the form 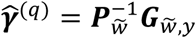, where 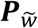 is the phenotypic (co)variance matrix of the first *q* PCs and 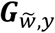 is a vector containing the genetic covariances between each of the top *q* PCs and the selection objective. Since the left-singular vectors are orthonormal (i.e., 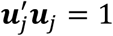 and 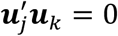, for *j* ≠ *k*), then 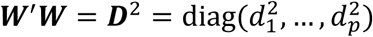. Hence, a method-of-moments estimate of the phenotypic (co)variance matrix of 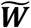 contains only the first *q* elements 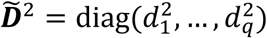; this is

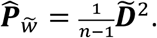

Using 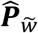 makes the coefficients of the PCs to be proportional to the genetic covariance between each of the PCs and the selection objective: 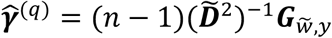. This solution can be mapped to coefficients for the measured traits using 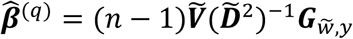, where 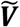 is the matrix containing only the first *q* right-singular vectors.

### Penalized selection indices

The objective function of the penalized SI is given by equation (3). Here we considered PSIs using either L1 or L2-norms or a combination of the two.

#### L2-PSI

Using an L2-norm as penalty, 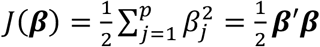, in equation (3) leads to the following optimization problem:

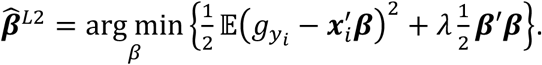

Therefore:

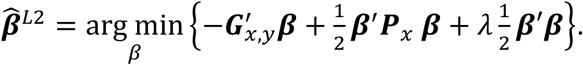

The second and third right-hand side terms can be combined to obtain:

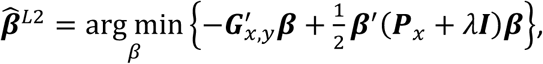

where ***I*** is a *p* × *p* identity matrix. Differentiating with respect to ***β*** and setting the derivatives equal to zero, we obtain the first-order conditions: 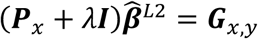; therefore:

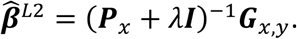

#### EN-PSI

The coefficients for the elastic-net family are obtained by considering an objective function as in equation (3), with 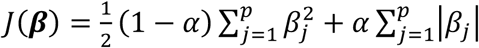; therefore,

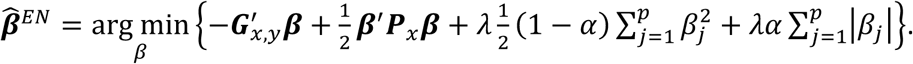

The L1-PSI and L2-PSI are particular cases corresponding to *α* = 1 and *α* = 0, respectively. When *α* = 0 the solution has a closed-form (see L2-PSI above). If *α* > 0, no closed-form solution exists; however, a solution can be obtained using the same iterative algorithms that are used to solve elastic-net regressions (e.g., LARS and coordinate descent^14^). These algorithms can be implemented either by ‘partial residuals’ or using ‘covariance updates’^32^. In our case, the objective function is entirely based on (co)variance terms. The objects ***P***_*x*_ and ***G***_*x,y*_ enter in the objective function of the PSI in the same way that ***X***′***X*** and ***X***′***y*** enter in a standard elastic-net regression. Therefore, to obtain solutions, we implemented the standard LARS algorithm (e.g., Hastie *et al*.^14^) entirely based on covariance updates.

### Data

The data set consists of 1,092 inbred wheat lines grouped into 39 trials and grown during the 2013-2014 season at the Norman Borlaug experimental research station in Ciudad Obregon, Sonora, Mexico. Each trial consisted of 28 breeding lines that were arranged in an alpha-lattice design with three replicates and six sub-blocks. The trials were grown in four different environments: *Flat-Drought* (sowing in flat with irrigation of 180 mm through drip system), *Bed-2IR* (sowing in bed with 2 irrigations approximately 250 mm), *Bed-EHeat* (bed sowing 30 days before optimal planting date with 5 normal irrigations approximately 500 mm), and *Bed-5IR* (bed sowing with 5 normal irrigations). In 2013, all the trials were planted by mid-November (optimal planting date), on the 21^st^ (*Bed-2IR* and *Bed-5IR*) and on the 26^th^ for *Flat-Drought*. Trials for *Bed-EHeat* were planted on October 30^th^. Grain yield (ton ha^−1^, total plot yield after maturity) was recorded.

Reflectance data were collected from the fields using both infrared (A600 series Infrared camera, FLIR, Wilsonville, OR) and hyper-spectral (A-series, Micro-Hyperspec, VNIR Headwall Photonics, Fitchburg, MA) cameras mounted on a Piper PA-16 Clipper aircraft on 9 different dates (time-points) between January 10^th^ and March 27^th^, 2014. During each flight, data from *p* = 250 wavebands ranging from 392 to 850 nm were collected for each pixel in the pictures. Using ArcMap software (ESRI, CA), the average reflectance of all the pixels within each geo-referenced trial plot was calculated and reported as a single data-point for each genotype for each band. Days to heading were recorded as the number of days from the date of sowing/first irrigation until 50% of spike emergence in each plot. Heading of about 50-80% of the total number of plots was used as criterion to distinguish between vegetative (VEG) and grain filling (GF) stages. The crop was considered to be at maturity (MAT) stage when the average RNDVI decreased to ∼0.4.

### Phenotype pre-processing

Within each environment, grain yield phenotypic records were pre-adjusted by fitting the following mixed model,

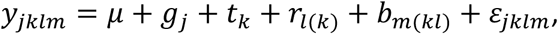

where *y*_*jklm*_ is the grain yield phenotype value for the *j*^th^ genotype, *k*^th^ trial, *l*^th^ replicate (within trial), *m*^th^ sub-block (within trial and replicate), *μ* is the overall mean and *g*_*j*_, *t*_*k*_, *r*_*l*(*k*)_, and *b*_*m*(*kl*)_ are the genotype, trial, replicate, and sub-block effects, respectively (all assumed to be random) and *ε*_*jklm*_ is an error term. Random effects were assumed to be independently and identically distributed (*iid*) normal with null mean and effect-specific variances. Likewise, the error terms were assumed to be *iid* with null mean and common error variance.

Grain yield data were pre-adjusted by subtracting from the phenotypic record (*y*_*jklm*_) the mean 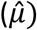 plus BLUPs of trial, replicate, and sub-block effects; this is

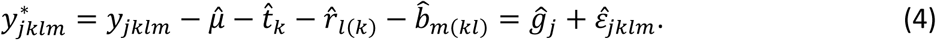

Reflectance data was pre-adjusted by fitting the above model, using reflectance at individual bands as phenotype expanded with the inclusion of a time-point effect. Separate models were fitted to each of the wavebands. As with grain yield, reflectance data were pre-adjusted by subtracting from the measured reflectance the estimated mean and predicted time-point, trial, replicate, and sub-block effects.

For quality control, pre-adjusted grain yield and reflectance phenotypes were removed for those grain yield scores lying beyond 3 times the inter-quantile region from the 0.25 and 0.75 quantiles.

After pre-adjusting, all phenotypes were standardized (to have unit variance); for ease of exposition, hereinafter we refer to the adjusted-scaled phenotypes (including grain yield and the image data) simply as phenotypes.

### Heritability estimation

After pre-adjusting standardization, we analyzed phenotypes using a mixed model of the form

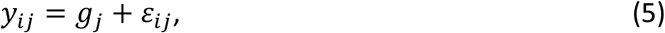

where *y*_*ij*_ is the phenotype for the *i*^th^ observation (*i* here is a single index for indices *k, l*, and *m* in equation (4)) of the *j*^th^ genotype; the genetic values are 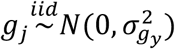, where 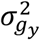 is the genetic variance; and the environmental terms are 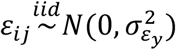. Plot-basis heritability was calculated from variance components estimates using

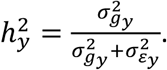

### Training-testing partitions

The data set contains information from 39 trials with 84 observations each. To assess the accuracy of indirect selection, we randomly assigned complete trials to testing sets. The training set comprised all the data from the trials not assigned to the testing set. This approach guarantees that no data from a single trial is present in both training and testing sets. This approach aims at representing a situation where one has calibrated the coefficients of the index using historical trials and apply these coefficients to image data of future trials. A similar validation scheme has been used (using herd-year-season groups instead of trials) in validation problems in previous studies involving milk spectra data^33^. In each training-testing partition, out of the 29 trials available, 26 trials (*n*_*trn*_ ≈ 2,184 observations) were randomly assigned to the training set, and the data from the remaining 13 trials (*n*_*tst*_ ≈ 1,092) was used for testing set. The regression coefficients of the indices (the ***β***’s for the canonical SI, PSI, and PC-SI) were calculated using grain yield and reflectance data of the training set. Estimates of the coefficients and reflectance data were then used to calculate the SI, 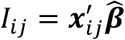, for each observation *i* in the testing set (*i* = 1, …, *n*_*tst*_). The heritability of the index and the genetic correlation between the index and the selection goal were estimated in the testing set.

The training-testing procedure described above was repeated 100 times by randomly assigning trials to training and testing sets. From these analyses, we reported the mean of heritability, genetic correlation, and selection accuracy; and their standard deviation across training-testing partitions.

### Estimation of phenotypic and genetic parameters

The population phenotypic (co)variance matrix ***P***_*x*_ was estimated within the training set using the unbiased sample (co)variance matrix given by 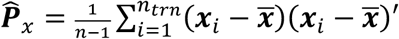, where 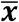 is the vector containing the sample mean of each waveband. Since reflectance data are centered and standardized, this reduces to 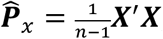, where ***X*** = [***x***_1_, ***x***_2_, …, ***x***_*n*_]′ is the matrix containing all measured traits in the training set.

The genetic covariance 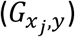 between grain yield and the *j*^th^ measured trait (*j* = 1, …, *p*) was estimated using a sequence of univariate genetic models as in equation (5). We fitted that model with grain yield phenotypes as response, then for each of the reflectance bands and then for the sum of grain yield and each of the bands. The genetic covariances between the bands and grain yield were then estimated using

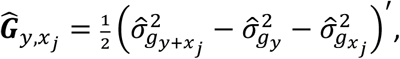

where 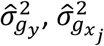 and 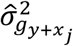 are the estimated genetic variances for grain yield, the *j*^th^ band, and the sum of grain yield and the *j*^th^ band, respectively.

### Estimation of the accuracy of indirect selection

To assess the accuracy of indirect selection we applied the regression coefficients derived in the training set to image data from the testing set to derive 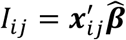. Then, using a mixed model analysis like that described in the previous section we estimated the heritability of the SI 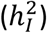, the heritability of grain yield 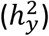, and the genetic correlation between the SI and grain yield 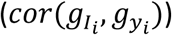. From these estimates, we derived the accuracy of indirect selection, 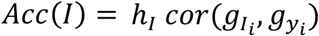, and the relative efficiency, 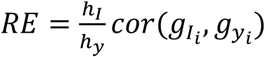.

### Software

All the aforementioned analyses were implemented in the R software environment^34^, version 3.5.1. Linear mixed models were implemented using the ‘lmer’ function from the LME4^35^ R-package. Code that implements LARS in the context of SIs was programed based on the LARS^36^ R-package.

### Materials and data availability

The data used in this study are publicly available by CIMMYT (https://www.cimmyt.org/) who owns all rights in the data. Data sets and R-scripts that perform all analyses are publicly available upon request to the corresponding author.

## Acknowledgments

We acknowledge CIMMYT’s Global Wheat Program that provided both experimental field and HTP data used in this work. M.L.C. was supported by the Monsanto’s Beachell-Borlaug International Scholarship Program (MBBISP).

## Author contributions

R.S., S.D., J.C. and S.M. were involved in the design of the field experiments and data collection. S.M. performed the HTP data correction and georeferencing. M.L.C. and G.D.L.C. conceived the idea, performed the analyses and produced a first draft. All the authors contributed to the final manuscript.

