## Supplementary Information for "Regularized selection indices for breeding value prediction using hyper-spectral image data"

This file contains:

- Supplementary Figs. S1-S8
- Supplementary Table S1

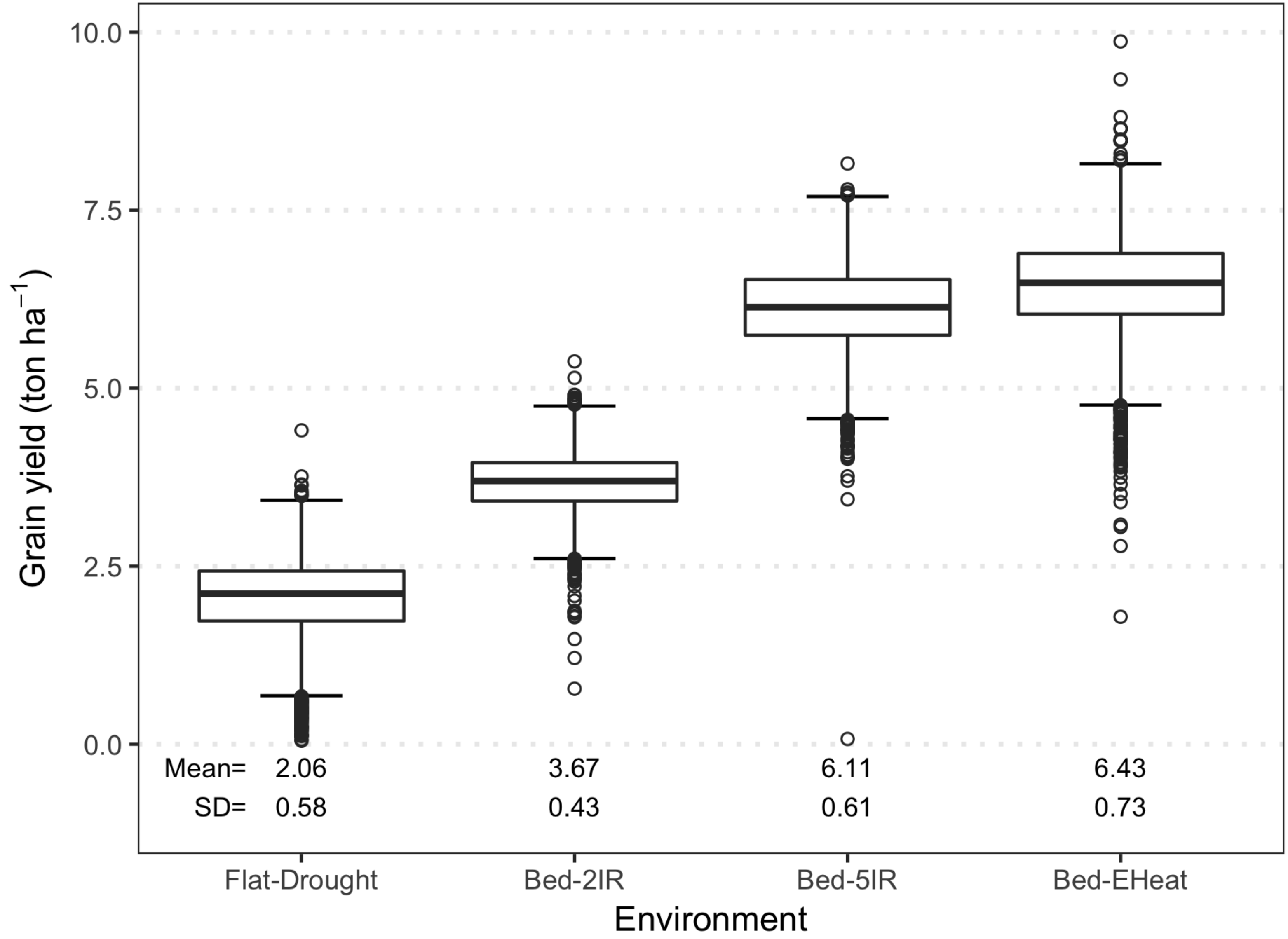


**Supplementary Fig. S1.** Box-plot of grain yield phenotypic records by environmental condition. $n \approx 3200$ observations within environment. SD: standard deviation.

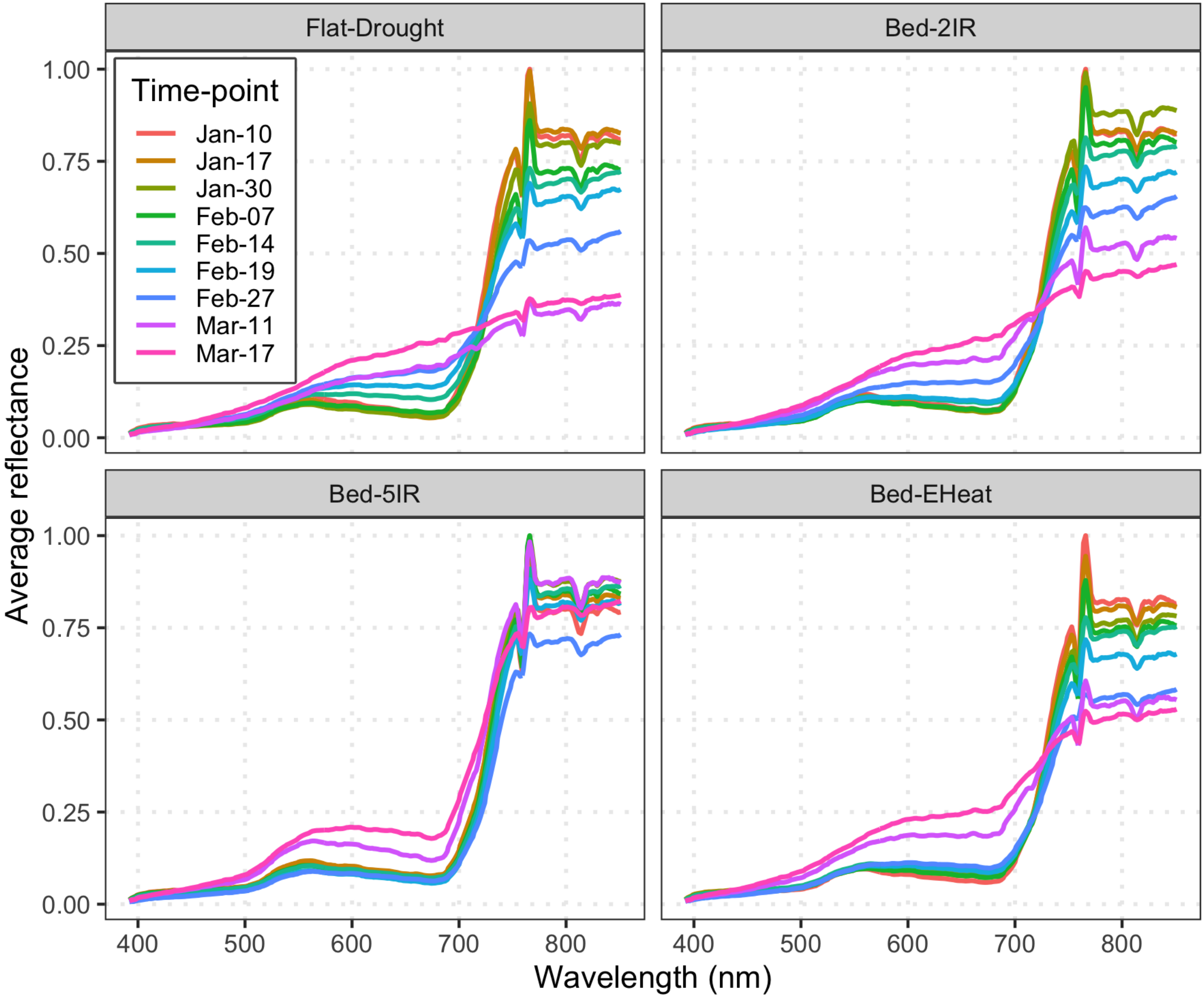


**Supplementary Fig. S2.** Light reflectance patterns as function of the wavelength. Each line represents the mean (across $n \approx 3200$ observations) reflectance for each waveband, within time-point (flight date). Within each environment, means were scaled to lie within 0 and 1 by dividing them by the maximum average.

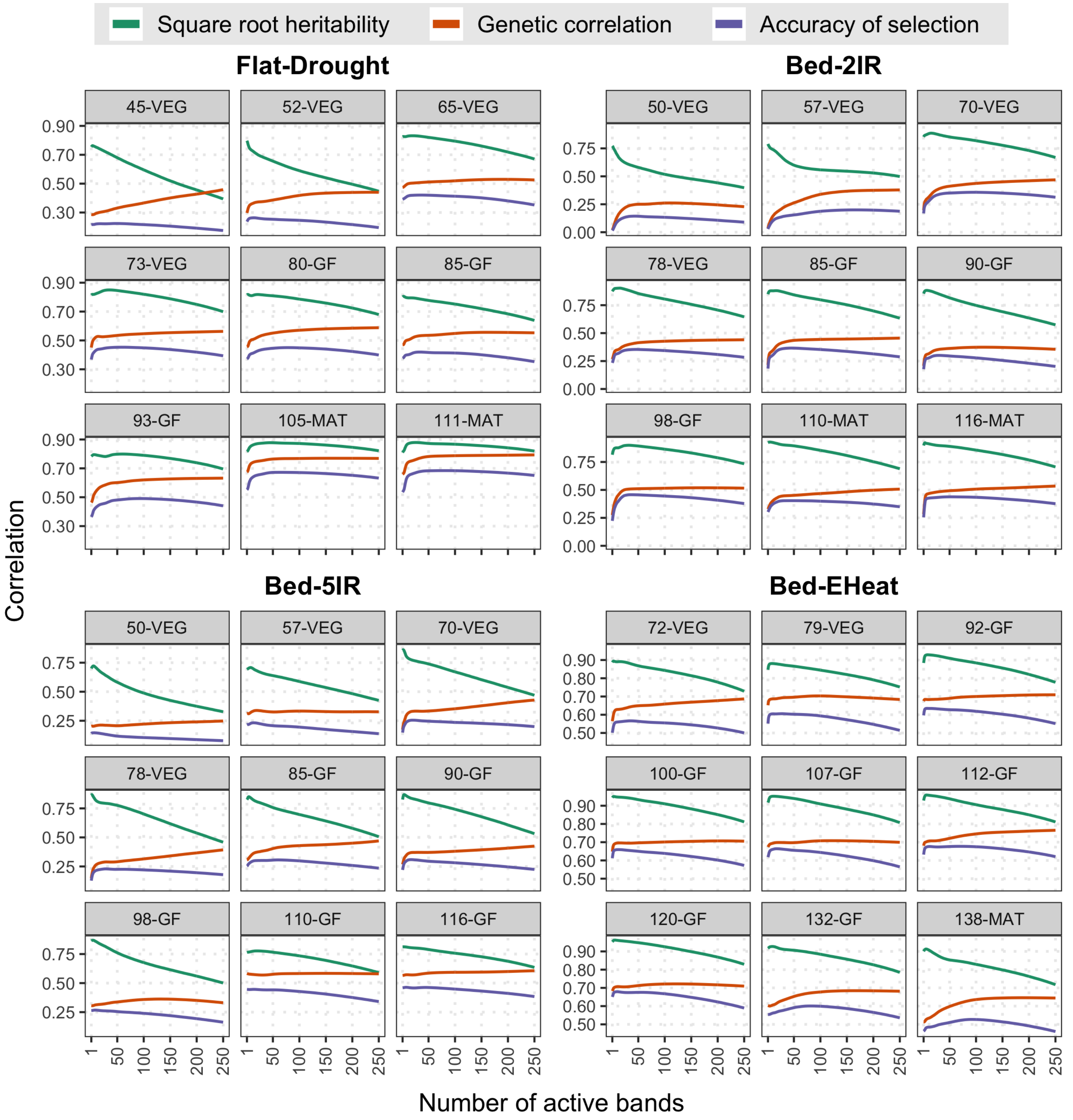


**Supplementary Fig. S3.** Accuracy of indirect selection of L1-PSI and its components. Square root heritability, genetic correlation and accuracy of indirect selection, all averaged over 100 training-testing partitions versus the number of bands entering in the index; by time-point (DAS=days after sowing, Stage: VEG=vegetative, GF=grain filling, or MAT=maturity) within environment.

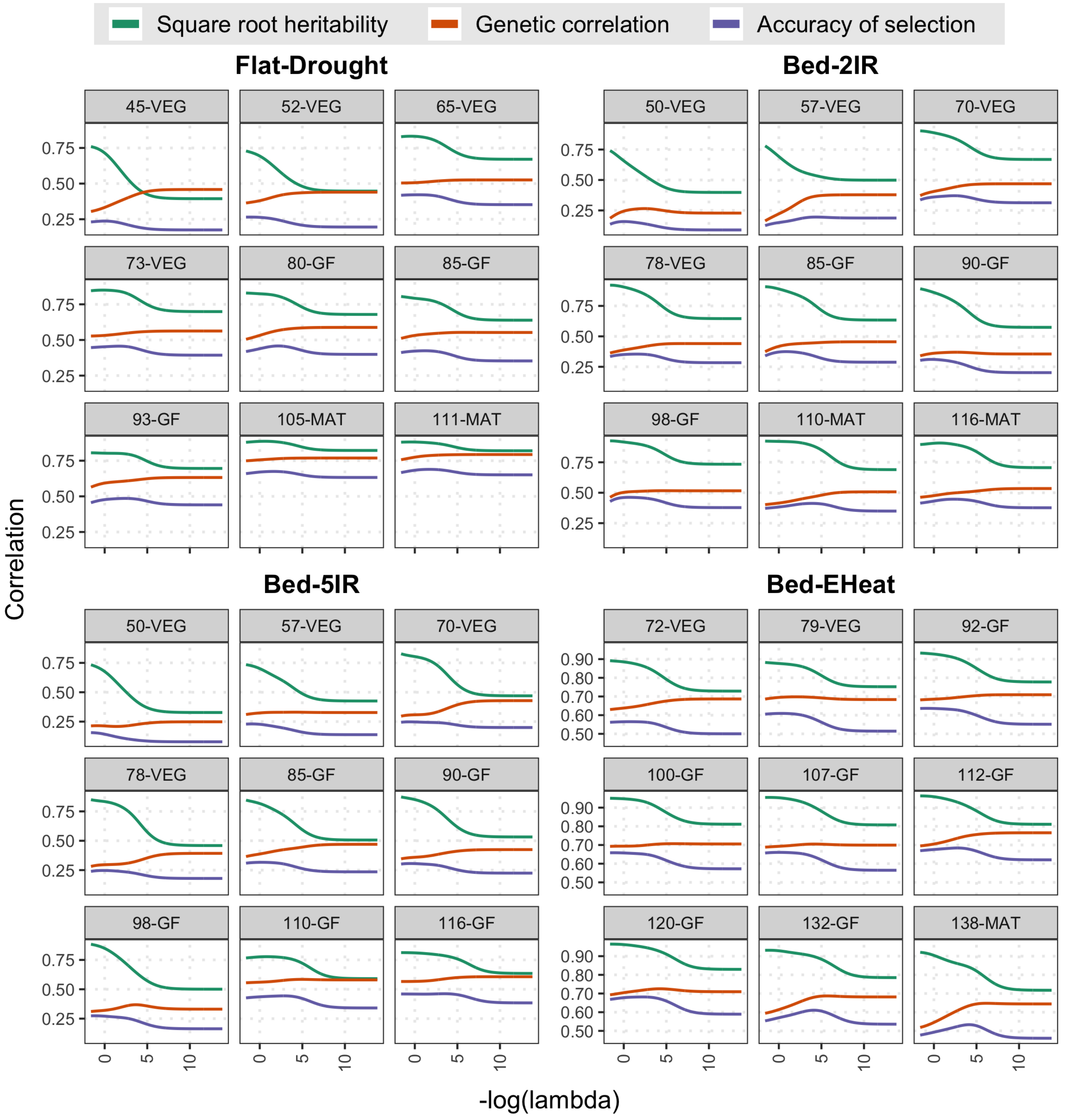


**Supplementary Fig. S4.** Accuracy of indirect selection of L2-PSI and its components. Square root heritability, genetic correlation and accuracy of indirect selection, all averaged over 100 training-testing partitions versus the penalization parameter ($\lambda$, logarithm scale) used to build the index; by time-point (DAS=days after sowing, Stage: VEG=vegetative, GF=grain filling, or MAT=maturity) within environment.

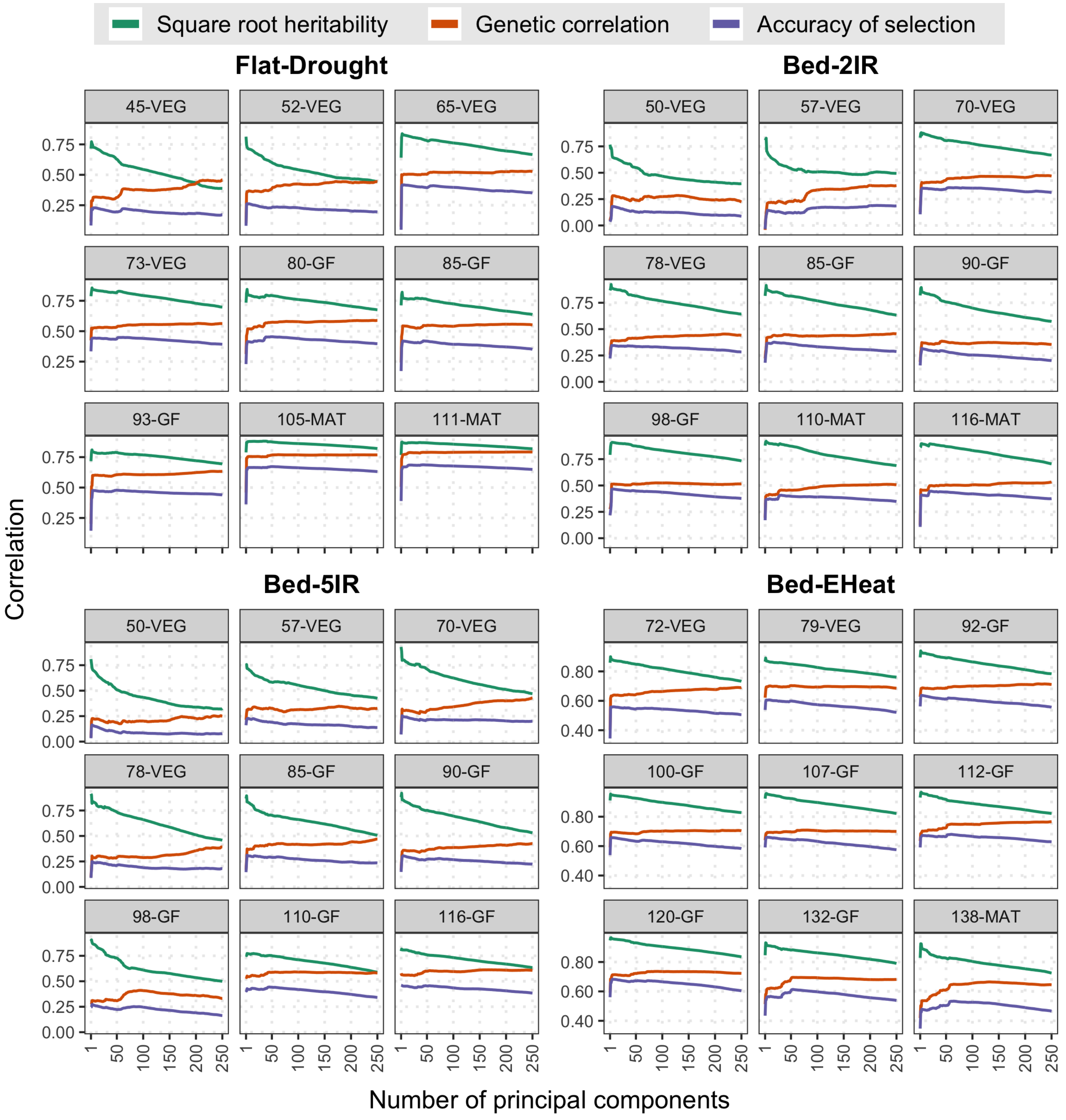


**Supplementary Fig. S5.** Accuracy of indirect selection of PC-SI and its components. Square root heritability, genetic correlation and accuracy of indirect selection, all averaged over 100 training-testing partitions versus the number of principal components used to build the index; by time-point (DAS=days after sowing, Stage: VEG=vegetative, GF=grain filling, or MAT=maturity) within environment.

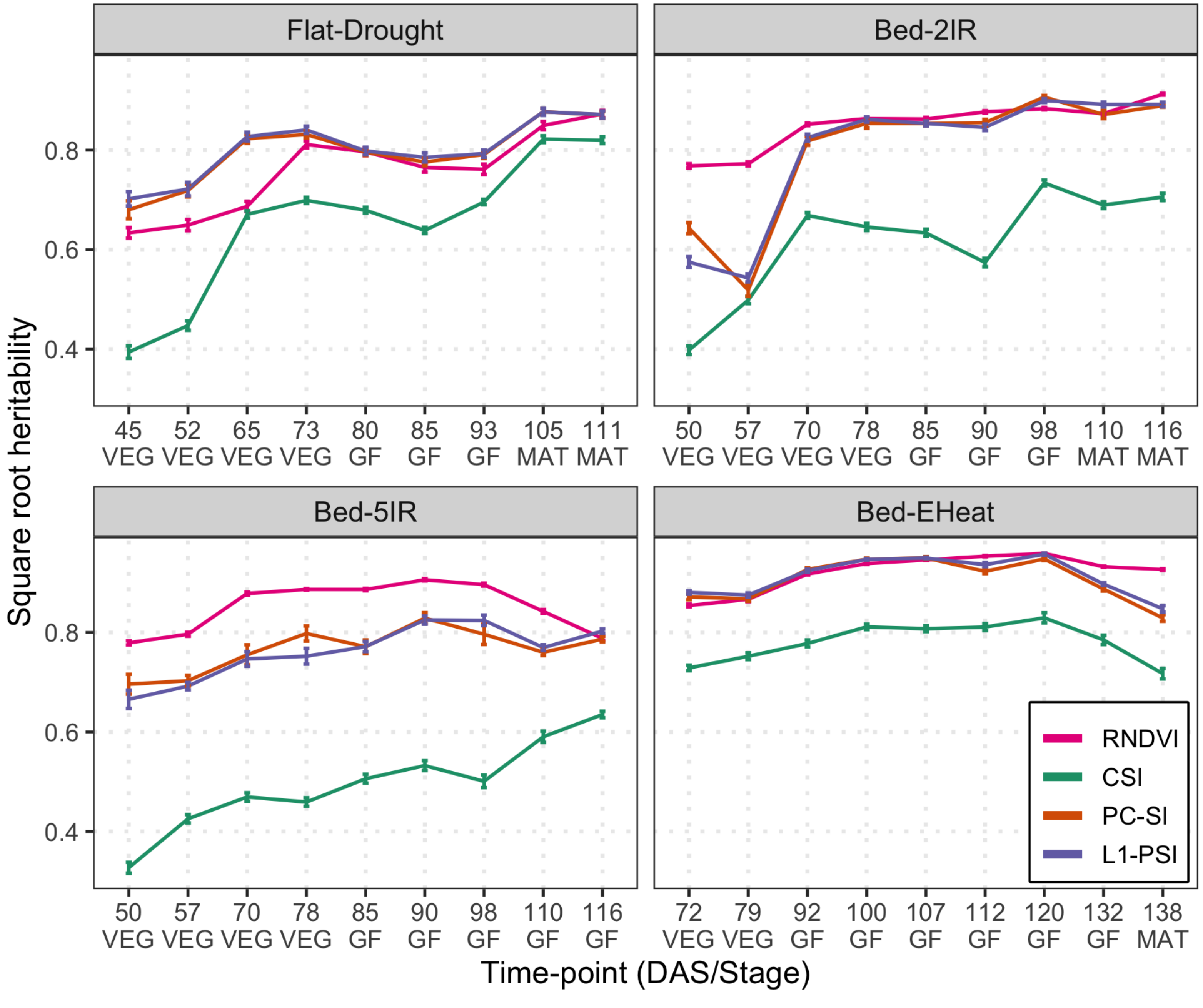


**Supplementary Fig. S6.** Square root of heritability of the canonical (CSI), of the regularized (PC-SI and L1-PSI) selection indices, and of the RNDVI. The lines provide the average square root heritability over 100 training-testing partitions. Vertical lines represent a 95% CI for the average. The horizontal axis give the time-point at which images were collected and are expressed in both days after sowing (DAS) and stages (VEG=vegetative, GF=grain filling, MAT=maturity).

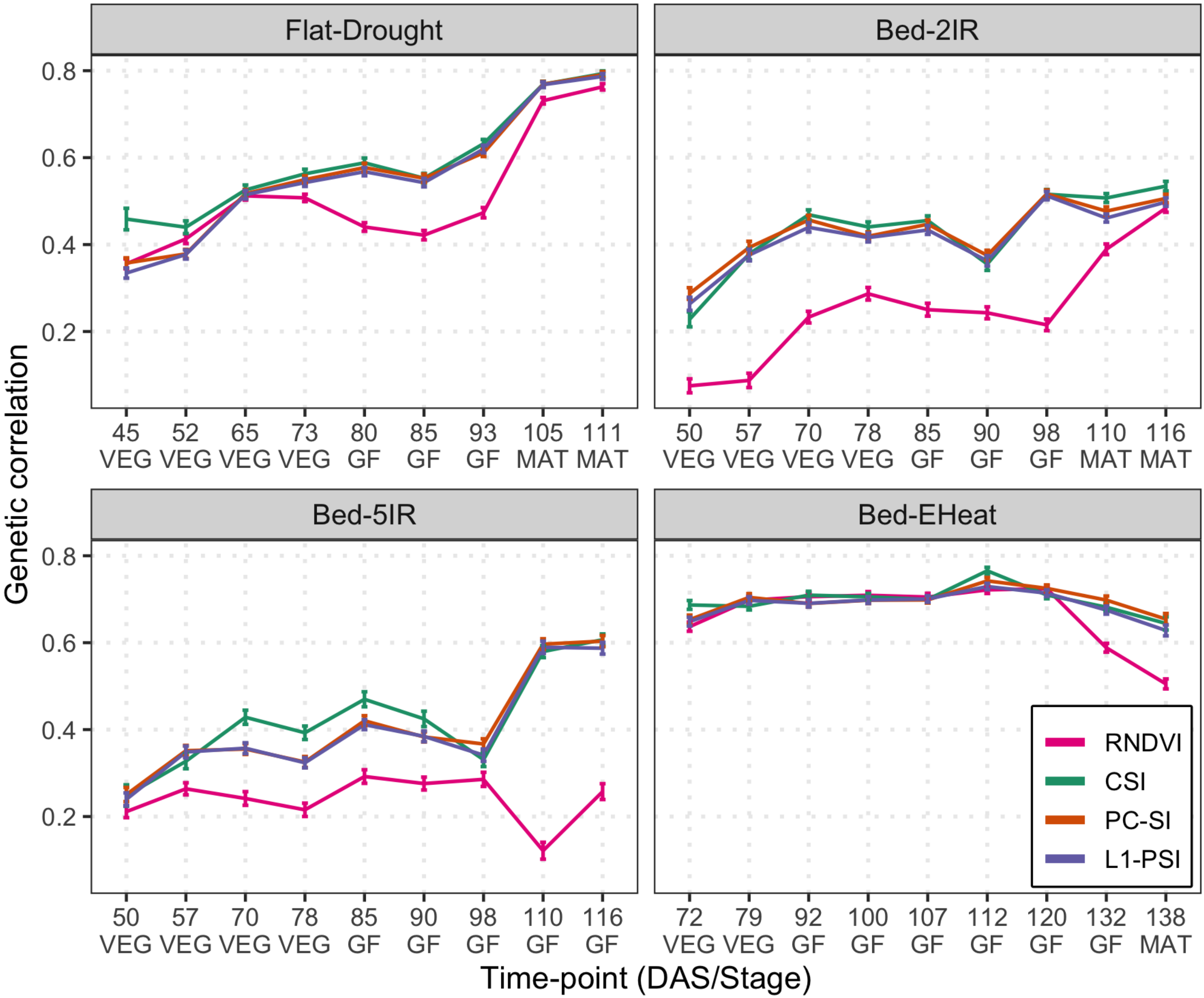


**Supplementary Fig. S7.** Genetic correlation between grain yield and all: the canonical (CSI), the regularized (PC-SI and L1-PSI) selection indices, and the RNDVI. The lines provide the average genetic correlation over 100 training-testing partitions. Vertical lines represent a 95% CI for the average. The horizontal axis give the time-point at which images were collected and are expressed in both days after sowing (DAS) and stages (VEG=vegetative, GF=grain filling, MAT=maturity).

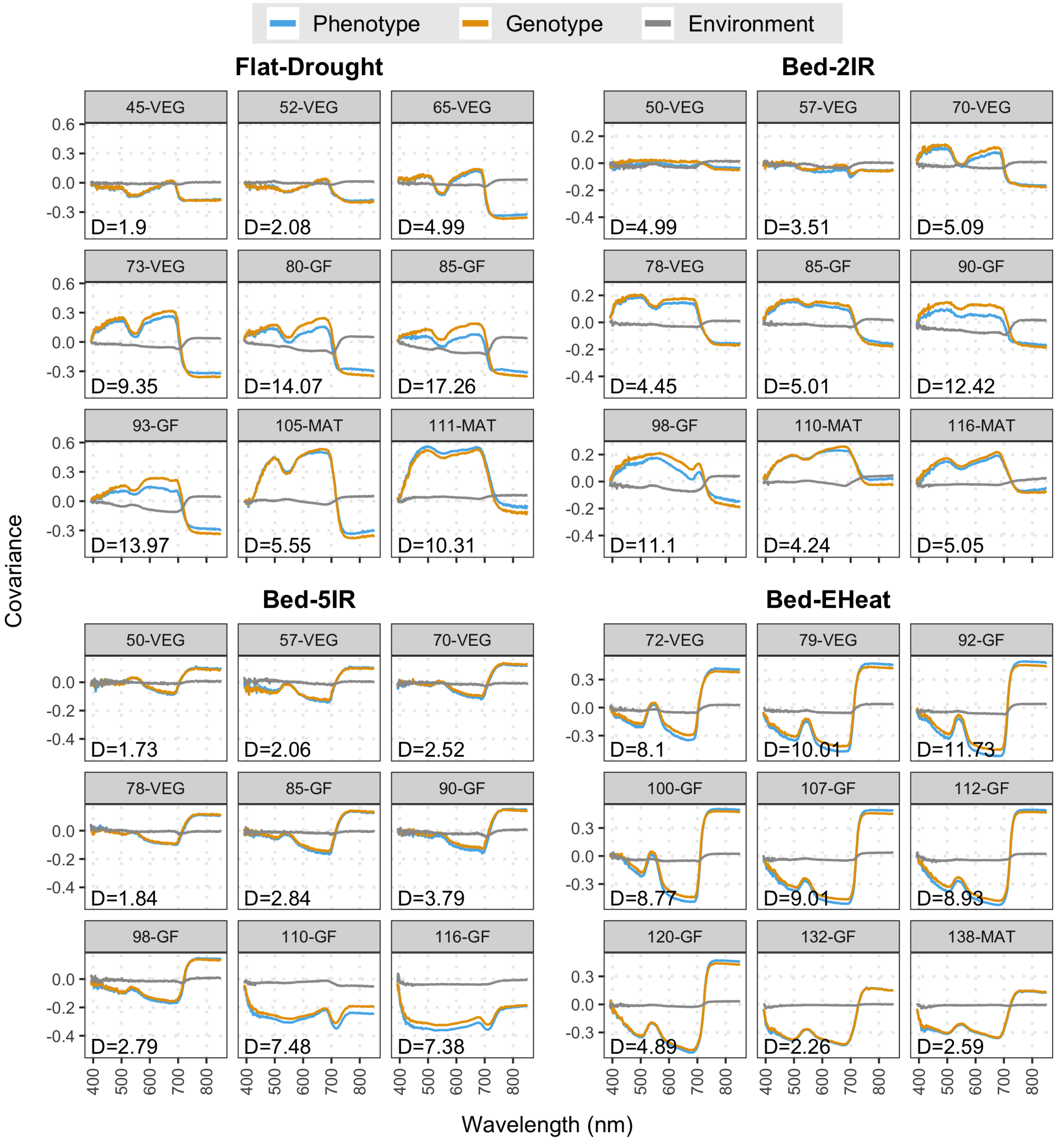


**Supplementary Fig. S8.** Phenotypic, genetic, and environmental covariances between wavebands and grain yield. 'D': discrepancy between phenotypic and genetic covariances as measured by the sum of the absolute differences; by time-point (DAS: days after sowing, Stage: VEG=vegetative, GF=grain filling, MAT=maturity) within environment.

| Env/time-point | | Phenotypic prediction | | | | Genotypic prediction | | | |
|---|---|---|---|---|---|---|---|---|---|
| | | PCR | L1-Phen | RNDVI | GNDVI | CSI | PC-SI | L1-PSI | L2-PSI |
| Flat-Drought | 45-VEG | 0.24 a | 0.23 a | 0.23 a | 0.21 b | 0.18 c | 0.24 a | 0.23 a | 0.24 a |
| | 52-VEG | 0.27 ab | 0.27 ab | 0.27 ab | 0.25 b | 0.20 c | 0.27 a | 0.27 a | 0.27 ab |
| | 65-VEG | 0.42 a | 0.42 a | 0.35 b | 0.35 b | 0.35 b | 0.43 a | 0.43 a | 0.42 a |
| | 73-VEG | 0.45 ab | 0.45 ab | 0.41 cd | 0.43 bc | 0.39 d | 0.46 a | 0.46 a | 0.46 a |
| | 80-GF | 0.44 bc | 0.43 c | 0.35 e | 0.39 d | 0.40 d | 0.46 a | 0.45 ab | 0.46 a |
| | 85-GF | 0.41 abc | 0.40 cd | 0.32 f | 0.39 d | 0.35 e | 0.43 a | 0.43 ab | 0.43 a |
| | 93-GF | 0.46 bc | 0.47 abc | 0.36 e | 0.45 cd | 0.44 d | 0.48 ab | 0.49 a | 0.49 a |
| | 105-MAT | 0.67 a | 0.67 a | 0.62 b | 0.64 b | 0.63 b | 0.68 a | 0.67 a | 0.68 a |
| | 111-MAT | 0.68 ab | 0.68 ab | 0.67 bc | 0.64 d | 0.65 cd | 0.69 ab | 0.69 ab | 0.69 a |
| | Multi | 0.68 cd | 0.68 bcd | 0.68 d | 0.65 e | 0.00 f | 0.70 ab | 0.70 abc | 0.70 a |
| Bed-2IR | 50-VEG | 0.18 a | 0.14 cd | 0.00 f | 0.12 d | 0.09 e | 0.18 a | 0.15 bc | 0.16 ab |
| | 57-VEG | 0.19 a | 0.19 a | 0.00 c | 0.03 b | 0.19 a | 0.20 a | 0.20 a | 0.20 a |
| | 70-VEG | 0.37 a | 0.36 a | 0.20 c | 0.21 c | 0.31 b | 0.37 a | 0.36 a | 0.38 a |
| | 78-VEG | 0.35 a | 0.35 a | 0.25 c | 0.30 b | 0.28 b | 0.36 a | 0.36 a | 0.36 a |
| | 85-GF | 0.37 a | 0.36 a | 0.22 c | 0.30 b | 0.29 b | 0.38 a | 0.37 a | 0.38 a |
| | 90-GF | 0.30 abcd | 0.29 cd | 0.21 e | 0.28 d | 0.20 e | 0.32 a | 0.31 abc | 0.32 ab |
| | 98-GF | 0.45 a | 0.46 a | 0.18 d | 0.35 c | 0.38 b | 0.47 a | 0.46 a | 0.46 a |
| | 110-MAT | 0.40 abc | 0.39 bc | 0.34 d | 0.39 c | 0.35 d | 0.42 a | 0.41 ab | 0.42 a |
| | 116-MAT | 0.44 a | 0.44 a | 0.44 a | 0.39 b | 0.38 b | 0.45 a | 0.44 a | 0.45 a |
| | Multi | 0.53 cd | 0.53 d | 0.46 e | 0.40 f | 0.01 g | 0.55 ab | 0.54 bc | 0.56 a |
| Bed-5IR | 50-VEG | 0.18 a | 0.17 ab | 0.16 ab | 0.15 b | 0.08 c | 0.17 ab | 0.16 ab | 0.16 ab |
| | 57-VEG | 0.25 a | 0.25 a | 0.21 c | 0.21 bc | 0.14 d | 0.25 a | 0.24 a | 0.24 ab |
| | 70-VEG | 0.27 a | 0.26 a | 0.21 b | 0.19 b | 0.20 b | 0.27 a | 0.27 a | 0.26 a |
| | 78-VEG | 0.26 a | 0.24 a | 0.19 b | 0.19 b | 0.18 b | 0.26 a | 0.24 a | 0.26 a |
| | 85-GF | 0.32 a | 0.32 a | 0.26 b | 0.25 b | 0.24 b | 0.32 a | 0.32 a | 0.33 a |
| | 90-GF | 0.31 a | 0.31 a | 0.25 c | 0.28 b | 0.22 d | 0.32 a | 0.32 a | 0.32 a |
| | 98-GF | 0.30 a | 0.29 a | 0.26 b | 0.25 b | 0.16 c | 0.29 a | 0.28 a | 0.28 a |
| | 110-GF | 0.46 a | 0.45 a | 0.10 d | 0.22 c | 0.34 b | 0.45 a | 0.45 a | 0.45 a |
| | 116-GF | 0.47 a | 0.47 a | 0.20 d | 0.34 c | 0.38 b | 0.47 a | 0.47 a | 0.47 a |
| | Multi | 0.54 a | 0.54 a | 0.32 c | 0.37 b | 0.00 d | 0.54 a | 0.55 a | 0.55 a |
| Bed-EHeat | 72-VEG | 0.57 a | 0.57 a | 0.54 b | 0.53 b | 0.50 c | 0.57 a | 0.57 a | 0.57 a |
| | 79-VEG | 0.61 a | 0.61 a | 0.60 a | 0.58 b | 0.51 c | 0.61 a | 0.61 a | 0.61 a |
| | 92-GF | 0.64 a | 0.64 a | 0.65 a | 0.63 a | 0.55 b | 0.64 a | 0.64 a | 0.64 a |
| | 100-GF | 0.66 a | 0.66 a | 0.67 a | 0.65 a | 0.57 b | 0.66 a | 0.66 a | 0.66 a |
| | 107-GF | 0.66 a | 0.66 a | 0.67 a | 0.66 a | 0.56 b | 0.66 a | 0.67 a | 0.66 a |
| | 112-GF | 0.68 ab | 0.68 ab | 0.69 a | 0.66 b | 0.62 c | 0.68 a | 0.68 a | 0.69 a |
| | 120-GF | 0.69 a | 0.68 a | 0.69 a | 0.66 b | 0.59 c | 0.69 a | 0.68 a | 0.68 a |
| | 132-GF | 0.62 a | 0.61 a | 0.55 b | 0.54 b | 0.54 b | 0.62 a | 0.61 a | 0.61 a |
| | 138-MAT | 0.54 a | 0.53 a | 0.47 b | 0.46 b | 0.46 b | 0.54 a | 0.53 a | 0.54 a |
| | Multi | 0.71 a | 0.70 a | 0.70 a | 0.67 b | 0.00 c | 0.71 a | 0.71 a | 0.72 a |

**Supplementary Table S1.** Accuracy of indirect selection (average over 100 training-testing partitions) for best phenotypic prediction (principal components (PCR), L1-penalized prediction (L1-Phen), RNDVI, and GNDVI) and for best genotypic prediction (canonical SI (CSI), optimal PC-SI, L1-PSI, and L2-PSI). Each row contains results for each environment and time-point (DAS: days after sowing, Stage: VEG= vegetative, GF=grain filling, MAT=maturity). Models with the same letter (within each row) are not significantly different from each other ($\alpha$=0.05, ANOVA followed by Tuckey test).
